# Endospore integrity and viability of *Bacillus subtilis* on simulated Martian regolith in rotational UVC radiation exposures

**DOI:** 10.1101/2025.09.19.677347

**Authors:** Gregory M Davis, Andrew R Greenhill, Jonathan Horner, Stephen C Marsden, Bradley D Carter

## Abstract

Current understanding of Martian regolith has advanced due to various rover explorations. Due to this, there are now several variants of Martian regolith that are chemically known and commercially available. These simulants are vital for panspermia models that suggest a transfer of simple life throughout the solar system via ejecta containing life from Mars when its surface was more favourable to host life. Bacteria that produce endospores are suitable candidates for these models as they have adequate protection from the harsh conditions of space, particularly against solar radiation. To assess this, the simple endospore former, *Bacillus subtilis*, was assessed for growth on several Martian regolith that represented different locations and epochs of Mars. This consisted of a conventional Martian regolith, a sulphur rich regolith, and a simulant of the Jezero crater which was thought to have once been flooded with water. We found that the sulphur rich regolith inhibited endospore formation, while the other two variants favoured endospore production. Interestingly, we also identified that pulsing of UVC in a simulation of endospores on rotational ejecta show that endospores break down faster with lower rotational frequency despite receiving the same UVC dose. Moreover, and most strikingly is that viable endospores after surviving UVC dosage displayed elevated expression of the DNA damage SOS response gene, RecA. Importantly, this study suggests that astrobiological approaches that utilise endospore viability as a benchmark for survival require reassessment as genomic integrity may be compromised.

**Importance:** Current panspermia models that explore the potential interplanetary transfer of simple microbes, particularly those that focus on Mars, apply a combination of Martian topology and bacterial survival. This study assesses *Bacillus subtilis* endospore formation on different Martain regolith mediums and simulates their potential exposure on fast and slow rotations while being exposed to UV radiation to simulate space radiation. We outline what regions of Mars would be suitable for the formation of endospores, and moreover, we assess viable endospores post-exposure for the expression of the SOS DNA damage repair gene, RecA. Endospores that are viable after withstanding near lethal doses of UVC show elevated expression of RecA. While our study adds vital information to understanding Martian topography and microbial survival, our finding RecA expression analysis post-exposure adds important information to our understanding of endospores as a whole and should be factored into future astrobiological approaches.

## Introduction

Our current understanding of Martian regolith and topography has been enhanced due to several missions that have explored the surface of Mars (reviewed in (1)) . The discovery of asteroids of Martian origin have also contributed vital information to our understanding of the chemical composition of Mars (2-4). As a result, we now have a strong understanding of the chemical composition of the Martian regolith across several regions of Mars’ surface. However, surface-based analysis of Martian regolith is based on current conditions, and does not necessarily accurately resemble when Mars had the potential means to sustain life (5-7). Nevertheless, these results mean that researchers attempting to model panspermia now have access to accurate compositions for the surface of different regions of the surface of Mars, which can therefore be simulated in a laboratory environment (8-10). When factoring in the conditions of space, and the potential viability of simple organisms to survive such conditions, these synthetic variants offer a unique way to explore the survivability of simple life forms that are delivered from Earth to land on an early Martian surface.

Panspermia models are based on the idea that life can be transferred from one planet to another through space, and is typically discussed in the context of the transfer of planetary ejecta carrying simple life forms through an interplanetary transfer (11-14). Bacteria that form endospores are considered to be an ideal candidate for such studies, which encapsulate bacteria with a protective wall that shelters bacterial DNA from harsh environmental conditions, including ultraviolet (UV) radiation (15, 16). Endosporulation has been intensively investigated in the bacterium *Bacillus subtilis*, making it an amendable model for astrobiological studies (11, 17-20). For this reason, *B. subtilis* endospores have been subjected to UV in the vacuum of space through the EXPOSE-E and EXPOSE-R missions in varying conditions (18, 21). Additionally, *B. subtilis* endospores have been exposed to the pressure they would sustain on ejecta to escape the surface of Mars (20). Therefore, the proposal that life was delivered to the Earth from Mars on asteroids that harboured simple life has been explored in various aspects.

Asteroids moving through space can be subjected to moderate or intense rotation, and factors including the size, and trajectory post-ejecta can influence rotational frequency (22-24). Larger asteroids usually display rotation periods of a few hours or longer (25, 26). However, smaller asteroids, of sizes comparable to the fragments that would be ejected from Earth or Mars as a result of an impact event, can display extremely rapid rotation (27, 28). In addition, the rotation rates of asteroids can change dramatically over time, as the result of non-gravitational forces driving both spin-up and spin-down processes (29-31). Therefore, the rotation of asteroids will clearly impact the way locations on the surface are exposed to UV radiation. Notwithstanding this, a rapid rotator and slow rotator would have the same UV exposure over a long period of time but would receive that exposure in a markedly different fashion. Rapid rotators will quickly alternate between illumination and dark, leading to the UV irradiation of the surface cycling rapidly. By contrast, the surface of a slow rotating asteroid will experience lengthy periods of irradiation followed by long periods of darkness. These variables can affect the time that the face of an asteroid is exposed to UV radiation in space, resulting in a shorter or longer time to reach a certain UV dose, influencing endospore survival.

We have previously shown that lethal UV dosage can be decreased if endospores are shielded by lysed bacteria on Martian regolith (11). In this study, different models of Martian regolith were used to determine endospore survival. We also simulate endospores on different regolith types for a variety of rotational frequencies, in order to assess the impact of the rotational nature of asteroids in space and demonstrate as a result that lethality is not simply dose dependant. Moreover, we show that endospores that were viable after exposure post-dosage of UV-C radiation show genomic expression of DNA damage repair mechanisms and suggest that endospore survival in these models are not correlative with bacterial genomic integrity.

## Materials and Methods

### Bacterial strains and generation of *B. subtilis* spores

This study utilised Strains *S. aureus* ATCC25923 and *B. subtilis* FUHAC10 (supplied by the Federation University culture collection). Strains were grown overnight in nutrient broth (NB) (ThermoFisher) with shaking at 37°C and growth was maintained on nutrient agar (NA) plates. Generation of *B. subtilis* spores were extracted using similar methods to those previously published (32). Briefly, *B. subtilis* vegetative cells were incubated by shaking at 37°C overnight in Schaeffer Sporulation Medium (SSM), then centrifuged at 10,000g for 10 minutes at room temperature before harvesting and the wash method described previously (33) was used for purification. Vegetative cells were destroyed by heat shock for 10 minutes at 80°C for 10 minutes then spores were stored in ddH20. Verification of spores Purity of spores was assessed using phase-contrast microscopy for purity and viability was verified by growing extracts on NA plates at 37°C overnight.

### Martian regolith acquisition and construction

Martian regolith simulants used in this study were purchased from Space Resource Technologies and compressed into small 35mm petri dishes prior to inoculation under sterile conditions. Simulants were as follows: MGS-1 Mars Glabal Simulant 003-05-001-1223, JEZ-1 Jezero Delta Simulant 003-08-001-1223. For the sulphur-rich simulant (SR), the MGS-1 simulant was used, supplemented with additional SO_3_ (Sigma-Aldrich) to reach the percentage as previously reported (9) which used the Paso Robles class soils as a reference. After addition of SO_3_, samples were mixed using a vortex prior to compaction.

### RNA extraction and cDNA synthesis

Post-exposure spores were carefully and aseptically collected using standard M9 buffer then transferred to sterile 1.5ml tubes. RNA isolation was achieved by using a TRIzol Max Bacterial Isolation Kit (Invitrogen). Samples were stored in RNase-free water and a TURBO DNA-free kit (Life Technologies) was used to remove any remaining DNA. Synthesis of cDNA was achieved as outlined in (34). Samples were suspended in RNAse-free water then used immediately for qRT-PCR analysis.

### Quantitative real-time PCR (qRT-PCR)

SYBR Green Taq Ready Jumpstart qPCR Mix (Sigma-Aldrich) was used for qRT-PCR. For each sample, the following was added: 5μL SYBR Green Mix, 0.5 μL of primers (2 μm), 0.15 μL of reference dye, and 10 μL of RNAse-free water. A no cDNA control and water controls were ran alongside each sample and all samples were run in triplicates on a Stratagene Mx300P (Agilent Technologies). Abundance was calculated using the using the ΔΔCt method (35) using the 16S rRNA gene as an internal standard. Primers used were as follows: RecA F: AACAGTTCGGCAAAGGTTCC RecA R: ATGAAGCGCCACAGTTGTTT 16S rRNA: F AGCAACGCGAAGAACCTTAC 16S rRNA R: ATTTGACGTCATCCCCACCT.

### UVC exposures

UVC radiation exposure was performed using an Opsytec BS-02 irradiation chamber and dosage was measured using an Opsytec UV-MAT dosage controller (Opsytec). For rotational exposures endospores placed on compressed regolith were rotated a battery operated RotoBo mini programmable rotator (Benchmark Scientific) inside the irradiation chamber and set for each RPM setting. Regolith was compressed into 20 small petri dishes (35mm in diameter) per exposure and prior to inoculation, spores were counted at 5.1 × 108 mL-1 (±1.0 × 108) then assessed by inoculating disks with 50μL droplets of purified spores prior to exposure. Each experiment was repeated three times, and the average of each exposure was calculated. Post exposure dishes were left to rest at room temperature for one hour before being transferred to NA plates and grown overnight at 37°C. Endospore survival and viability was assessed as growth post exposure per sample rather than at a cellular basis.

### Statistical analysis and software

The Prism 5 software package (Graphpad) was used to generate graphs and to perform statistical analysis. Statistical significance was performed using student t-tests and calculating standard deviation from the mean. Figures were processed and annotated using Adobe Illustrator (Adobe).

## Results

### Endospore survival on different Martain regolith

Endospore forming bacteria have previously been shown to subsist on Martian regolith in different representations of Martian environment. To expand on this, different simulants as described previously (8) were used to assess survival of spore forming and non-spore forming bacteria (Table 1). Collectively, these mediums represent different epochs of Martian regolith and environment and consisted of the following: The Jezero Delta Simulant (JEZ-1), based on samples collected by the Perseverance rover. The Jezero delta represents an ancient riverbed and has been explored for passible past signs of microbial life (36). For a contemporary regolith, the Mars Global Simulant (MGS-1) was used (8). MGS-1 is based on the Rocknest crater collected by the Curiosity rover (37), and although it does not represent a specific epoch of Mars, it represents a contemporary Martian regolith. Lastly, a Sulphur-Rich medium (SR) was constructed using the MGS-1 regolith supplemented with SO_3_. The sulphur enrichment was based on the SR media as described previously (9), which was from the Paso Robles class soils samples collected by the Spirit rover which are thought to have formed from hydrothermal condensates due to degassing of magma (38, 39). Broadly, this selection of Martian regolith gives a greater variance to assess endosporulation than observed in our previous study (11).

**Table 1:** Survival of bacteria on different regolith post-incubation.

| Regolith | *B. subtilis* spores | *S. aureus* |
| --- | --- | --- |
| JEZ-1 | Yes | No |
| MGS-1 | Yes | No |
| MGS-1(S) | No | No |

To assess whether bacteria could form endospores when exposed to each model regolith, liquid cultures of both the spore former, *B. subtilis*, and a non-spore former, *S. aureus* were inoculated onto small disks of compressed regolith then incubated at 37°C for 24 hours. After inoculation each disk was then analysed for the formation of endospores via microscopy then transferred to nutrient growth media to assess survival in the absence of spores (Table 1). As expected, *S. aureus* was unable to grow in the presence of either regolith and *B. subtilis* spores were observed for JEZ-1 and MGS-1 regolith variants. Surprisingly, the SR media was inhibitive for the generation of *B. subtilis* endospores. The optimal temperature for endosporulation in *B. subtilis* has been reported to range from 30°C to 45°C (40). Therefore, to rule out the possibility that the selected temperature of 37°C was not supportive of endosporulation, this was repeated at 30°C. 35°C, 40°C and 45°C (Table S1). Despite this temperature variance, endosporulation was unable to be achieved on the SR regolith, which therefore seems likely too toxic for the bacteria to generate endospores and were unable to survive. This suggestion is not purely speculative as there have been several studies that show an inhibition of endosporulation when *B. subtilis* is grown on a sulphide enriched medium (41, 42), which is due to the generation of sulphide which occurs under stress conditions. Therefore, it is possible that the high sulphur content in the SR medium inhibited the generation of endospores and hence, why no spores were observed across a broad temperature range. From here on, JEZ-1 and MGS-1 were used as candidate mediums for additional analysis.

### Pulsation of UVC irradiation affects endospore viability

To simulate the effects of UVC radiation in space, endospores were generated as a slurry, inoculated onto JEZ-1 and MBS-1 regolith mediums, then irradiated at varying doses (Figure 1A). Since both regolith mediums support the formation of endospores, the results upon exposure were similar to UVC exposures reported previously as expected (11, 32). Despite a minor variance where the JEZ-1 regolith consistently showed a higher death rate of spores on regolith disks than MGS-1, both mediums showed a consistent trend which is similar to that previously observed on nutrient growth media (11).

**Figure 1:**
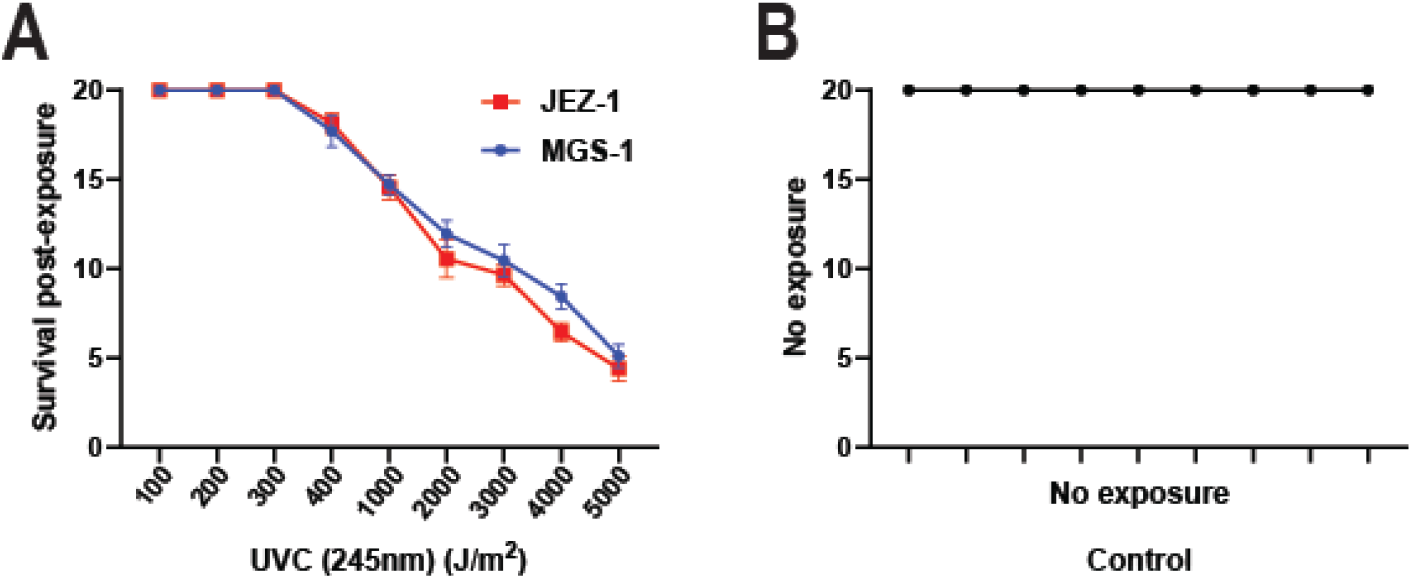
Endospore viability post-exposure to UVC radiation. **(A)** Vegetative *B. subtilis* cells were inoculated on disks of JEZ-1 (red squares) and MGS-1 regolith (blue circles), then subjected with UV-C radiation and confirmed viable by the presence of endospores **(B)** Control disks where vegetative *B. subtilis* cells were inoculated onto JEZ-1 and MGS-1 disks but did not receive UVC radiation. Error bars = SDM, N = 60.

Previous simulations have shown that meteorites of Martian ejecta would be subjected to pressure that endospore forming bacteria could survive (20). Therefore, to investigate the possibility of enhanced survival of endospores on simulated debris while rotating upon exposure to UVC, endospores on JEZ-1 and MGS-1 regolith disks were inoculated, then rotated at different frequencies. In this case, endospores were all irradiated with the same dosage of UVC, however, the frequency of rotation increased with each assay to simulate a pulsation like effect to model the day-night cycle upon a spinning object (Figure 2A, B and S1A). Interestingly, the same dosage given over slower rotational frequency had a greater detrimental effect on endospore viability than when disks were rotated at greater frequency. It should be noted that the dosage was consistent as the time of exposure for each rotational speed was calculated to give the same dose regardless of rotational frequency. This is peculiar, as varying results from previous studies have shown that pulsed delivery of UVC has been as effective with endospore exposure to constant dosage (43, 44). In addition to this, for non-spore forming bacteria, pulsating UVC has shown to be equal or greater in efficacy on sterilising bacterial growth (43-47). Nonetheless, this result clearly shows that a greater frequency of pulsation favours the longevity of endospores. This suggests that the penetrance of UVC is less effective with shorter, albeit more frequent exposure to the same UVC dose compared to those with less frequency.

**Figure 2:**
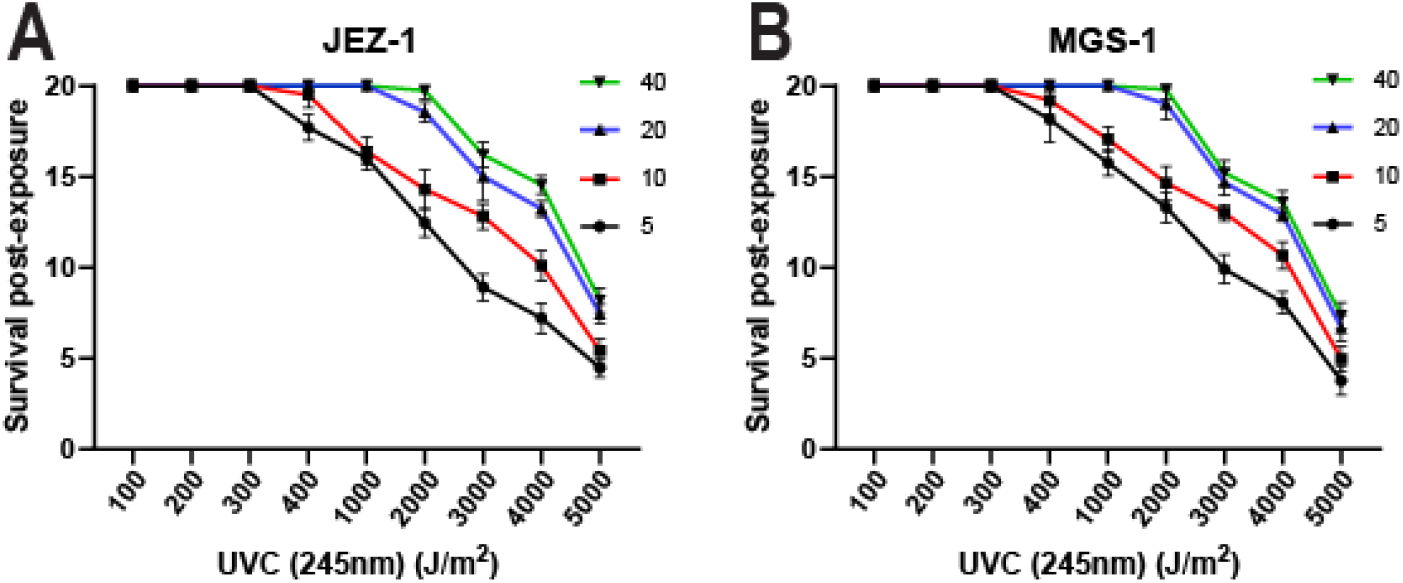
Intermittent UVC dosage in simulated rotational exposure assays: Endospores inoculated on disks of EB **(A)** and MGS-1 **(B)** and subjected to UVC radiation at different RPM frequencies at the same UVC dose. No exposure control (Figure S1A). RPM was as follows: black = 5RPM, red = 10RPM, blue = 20RPM & green = 40RPM. Error bars = SDM, N = 60.

### Viable endospores post-exposure exhibit signs of DNA damage repair

Exposure to UV radiation to double stranded DNA induces bonds between neighbouring pyrimidine groups which interfere with transcription (48, 49). This ultimately results in deleterious mutations or cell death. In bacteria, the SOS response is responsible for triggering a response to DNA damage (50, 51). This involves upregulation of the RecA gene which cleaves and activates the LecA SOS repressor, resulting in the upregulation of DNA damage repair genes. Due to this, upregulation of RecA is an ideal biomarker for the DNA damage response in bacteria (52-54). The survival of endospores is a critical component of many astrobiology models that involve panspermia. To assess the genomic integrity of endospores that were viable, the pulsing UVC exposures were repeated and the remaining disks with viable spores after a dosage of 5000 J/m^2^ were measured for RecA expression using qRT-PCR (Figure 3 and S1B). Strikingly, we observed higher values of RecA expression in endospores that were viable post-dosage. Moreover, the trend observed when UVC was delivered in a pulsation manner while spores were rotating was also evident with RecA expression. In all rotational frequencies there was a significant difference from controls of unexposed spores and unexposed vegetative cells.

**Figure 3:**
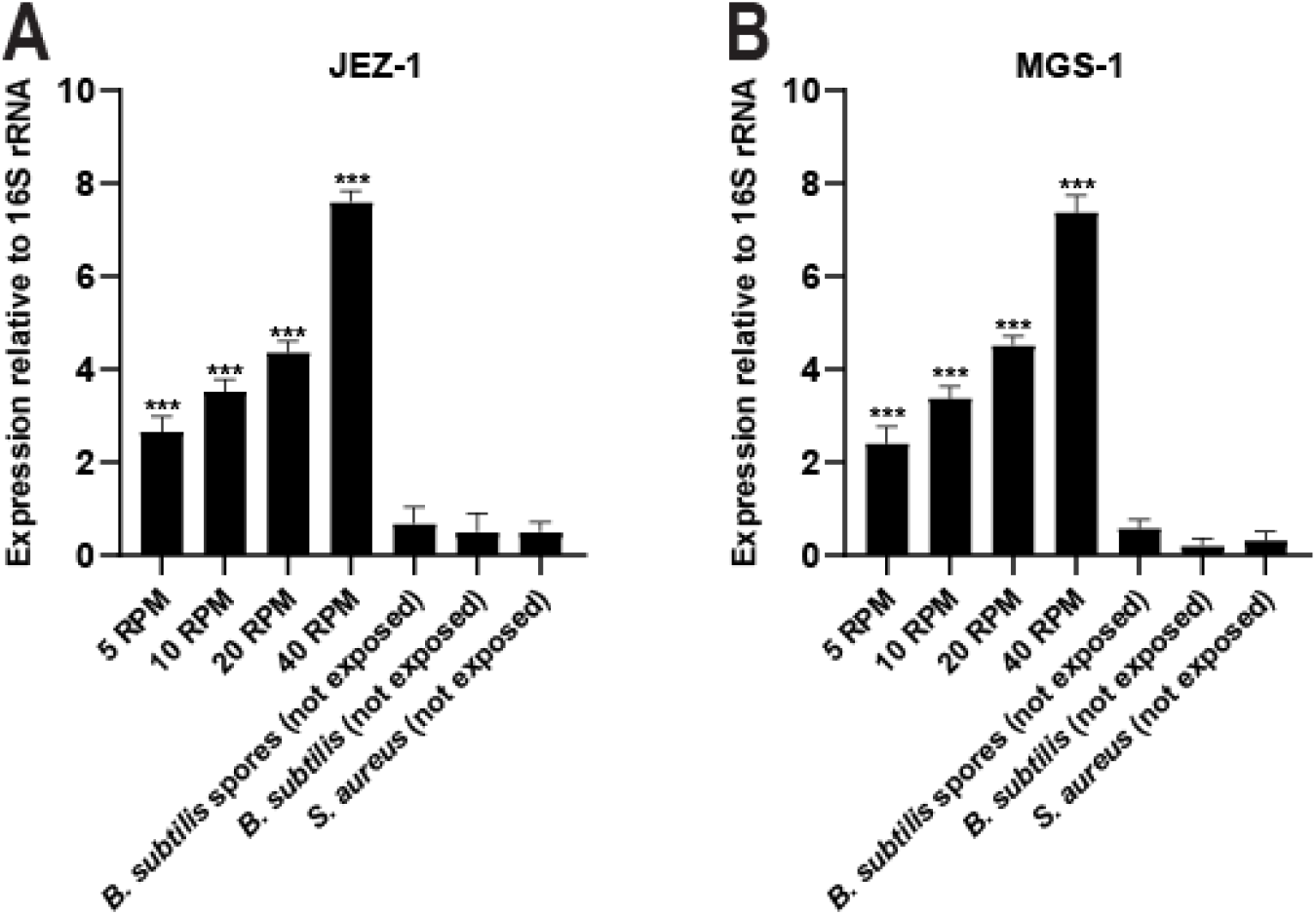
Viable spores display DNA damage response in viable endospores: qRT-PCR expression of RecA in viable spores recovered from UVC rotational exposures (5-40RPM) on **(A)** JEZ-1 and **(B)** MGS-1 regolith. Expression relative to 16S rRNA using no exposure control spores, vegetative cells, and the non-spore former *S. aureus* as controls. Error bars = SDM, N = 20, *** P ≤ 0.001.

To eliminate the possibility that the JEZ-1 or MGS-1 regolith had any impact on the integrity of *B. subtilis* endospores, this process was replicated without using Martian regolith. To achieve this, a slurry of endospores was placed on glass slides and exposed to both pulsing and non-pulsing UVC to a of dosage of 5000 J/m^2^ (Figure S1B). Spore viability was assessed via microscopy then subjected to the same qRT-PCR analysis of RecA expression. These results were consistent with those performed on EB and MGS-1 regolith, suggesting that both forms of Martian regolith exhibit no toxicity towards the formation of *B. subtilis* endospores. Moreover, this result suggests that using endospore survival and viability as an indicator of bacterial survival or robustness in panspermia models should be treated with caution as these endospores may be genomically compromised.

## Discussion

### Martian topography and endospores in simulated space radiation (UVC)

Panspermia models that include Martian ejecta have assessed numerous variables using endospores as their candidates for withstanding the initial impact, the pressure they would endure upon exiting Mars, and their survival in space (11, 13, 14, 17-21, 55). *B. subtilis* is an ideal candidate for these studies as it is arguably the most studied endospore former to date and can form endospores upon exposure to considerable toxic surfaces or when nutrient uptake is restricted (33, 56). In this study we have exposed vegetative *B. subtilis* cells to synthetic Martian regolith variants that represent different regions and epochs of Mars topography. Of the variants assessed, almost all were suitable for allowing the formation of endospores. Unexpectedly, the sulphur rich regolith that represents Paso Robles class soils inhibited the formation of endospores as none were viable after their inoculated on the SR media.

Although this was unexpected, there are potential reasons why a sulphur rich environment inhibits the time taken to form endospores, ultimately killing the vegetative cells. Sulphide has been shown to inhibit the formation of endospores, mostly as intracellular sulphur production occurs in *B. subtilis* as a stress response (41). This is associated with *B. subtilis* normally undergoing a desulphurisation process when metabolising intracellular cystine (a sulphur-containing amino acid) which gets converted to sulphide endogenously, resulting in ‘cysteine toxicity’ (42). Therefore, is it plausible that the excessive sulphur present in the SR medium may have resulted in toxicity, inhibiting the formation of endospores. Furthermore, it should be noted that the limitation of all Martian regolith models is that they are all based on either meteorite samples of Martian ejecta, or recent collections of rover-based soil samples from Mars. This places significant limitations on their use for assessing bacterial survival, as none of these candidates are a realistic regolith of Mars when it potentially had conditions that could harbour life. Therefore, these results need to factor in that they resemble the conditions of the Martian surface now, which are much less likely to accommodate life than those found earlier (57).

Another variable that outlines a degree of caution when assessing current panspermia models is the variance regarding the penetrance of UVC radiation when delivered in a pulsating manner. Our results show that more frequent rotations of simulated Martian ejecta decrease the death rate of *B. subtilis* endospores. It is understood that larger asteroids rotate at slower speeds compared to smaller ones (22, 23). However, small asteroids and meteoroids can be expected to display a wide range of rotation rates - with asteroids in the metre-to-ten-metre size range known to exhibit rotation periods as fast as just a few seconds (58). In this light, our results suggest that small ejecta rotating at such high speeds could prove ideal vehicles for the transfer of viable material between planets, as the rapid rotation would result in a significant delay in the detrimental effects of radiation from space. While several studies have shown conflicting results regarding pulsating vs constant delivery of UVC radiation (43-47), less is known regarding endospore formers in this context. The increased frequency in pulsation in our results suggests that UVC has less penetrance than when rotating rapidly, despite the same dosage being received. Although, whether this delays the onset of DNA damage remains unknown in our observations.

### Endospore viability is not an accurate means to assess long term survival of *b. subtili*

Endospores form under times of stress, including lack of nutrients or toxicity in the external environment (59). Here, we show for the first time that the initial biomarker for initiation of the SOS response to DNA damage is upregulated in endospores exposed to UVC radiation. This finding is significant, not only as it adds new understanding to the microbiology of endospores, but also with the context of astrobiological modelling. Several panspermia models using endospores factor in endospore viability as an indicator of survival. However, these models should potentially be revised as upregulation of RecA was observed in endospores that did not degrade in the irradiation process. Therefore, these bacteria potentially harbour mutations which might be detrimental to their function, suggesting that viability is not accurate as the genomes of these bacteria may be compromised.

To counter this as a problem, it could be argued that the survival of these bacteria, albeit it, with suspect mutations might be associated with driving evolution. This is particularly of interest in theories that suggest that early life may have arrived on Earth rather than have formed through an abiogenesis process. The earliest fossils so far discovered on Earth date back approximately 3.5 Gyr (60, 61). However, due to these early fossils, it is often suggested that life on Earth became established as soon as the planet was physically capable of supporting life. Prior to about 3.7 or 3.8 Gya, it is thought that the impact rate experienced by Earth was too high for life to become established. either because of the proposed Late

Heavy Bombardment (62, 63), or as the tail end of a smoother decay in the impact rate during the clean-up phase of the planet formation process (for detailed review, see (64)). Nonetheless, our study adds an interesting layer to the plausibility of this and other studies and suggests a degree of caution that needs to be considered when using endospore viability as a marker of bacterial survival and longevity.

## Acknowledgements

We acknowledge the technical staff at Federation University Australia for the donation of the bacterial strains used in this study. We also acknowledge the technical staff at Swinburne University for their assistance with the UVC exposure assays.

